# Reactive Oxygen Species Drive the Aberrant Immune Response to a *C. neoformans* Chitin Synthase 3 (*chs3Δ*) Mutant

**DOI:** 10.1101/2025.06.24.661434

**Authors:** Rebekah G. Watson, Camaron R. Hole

## Abstract

*Cryptococcus neoformans* is a globally distributed fungal pathogen that causes severe opportunistic infections, particularly in individuals with impaired cell-mediated immunity, such as AIDS patients and organ transplant recipients. Chitosan, a deacetylated derivative of chitin, plays a key role in modulating immune responses during *Cryptococcus neoformans* infection. We previously showed that mice inoculated with a heat-killed (HK) chitosan-deficient *C. neoformans* strain lacking chitin synthase 3 (*chs3Δ*) undergo rapid, neutrophil-dependent mortality within 36 hours, despite the absence of viable organisms. Here, we investigated the immune mechanisms underlying this lethal inflammatory response. We assessed the role of inflammatory cytokines, adaptive immunity, and neutrophil-derived effector functions in this lethal response. IL-6–deficient and Rag1–deficient mice exhibited only modest survival benefits, suggesting that IL-6 signaling and adaptive immune cells play limited roles. In contrast, mice lacking NADPH oxidase 2 (NOX2), the catalytic subunit of the phagocyte oxidase complex required for reactive oxygen species (ROS) production, were completely protected. Despite equivalent pulmonary neutrophil recruitment in wild-type and NOX2-deficient mice, only the former displayed elevated proinflammatory cytokine and chemokine levels and succumbed to disease. These findings indicate that neutrophils are not intrinsically pathogenic but mediate lethal immunopathology through ROS-dependent inflammatory amplification.

## INTRODUCTION

*Cryptococcus neoformans* is a globally distributed, encapsulated fungal pathogen that causes life-threatening pneumonia and meningoencephalitis, particularly in immunocompromised individuals. The most common risk group comprises people with impaired CD4⁺ T cell function, such as those with advanced HIV infection, where cryptococcal meningitis accounts for an estimated 19% of all AIDS-related deaths globally [1; 2]. Recent analyses estimate approximately 194,000 incident cases and 147,000 deaths per year due to cryptococcal meningitis, underscoring the urgent need for improved diagnostics, treatments, and preventive strategies [3]. Additional at-risk groups include solid organ transplant recipients, individuals receiving immunosuppressive therapy, and patients undergoing chemotherapy [4]. Despite the availability of antifungal drugs, treatment is prolonged, associated with high toxicity, and often inaccessible in low-resource settings [5]. Moreover, no licensed vaccine exists to prevent cryptococcal disease [6]. Thus, a deeper understanding of the immune mechanisms that mediate protection versus pathology in cryptococcosis is essential to guide the development of more effective therapeutic and preventive strategies.

A key immune evasion strategy employed by *C. neoformans* involves modification of its cell wall with chitosan, a deacetylated form of chitin that dampens immune activation and supports persistence in the host [7; 8; 9; 10]. Chitosan-deficient strains lacking chitin deacetylases (*cda1Δcda2Δcda3Δ*) are rapidly cleared from the lung and can confer protective immunity when delivered in a heat-killed form, serving as effective whole-cell vaccine candidates [11; 12]. In contrast, the *chs3Δ* mutant, which is also chitosan-deficient due to the deletion of chitin synthase 3, elicits a dramatically different outcome.

Inoculation with heat-killed (HK) *chs3Δ* yeast results in uniform, rapid mortality within 36 hours, accompanied by extensive neutrophilic infiltration and a pronounced cytokine storm involving IL-6, G-CSF, and CXCL1/KC [13]. Strikingly, this pathology occurs even in the absence of viable fungal cells, indicating that the host immune response is the primary driver of mortality in this model.

Our prior work established that neutrophils are central mediators of the rapid mortality induced by heat-killed (HK) *Cryptococcus neoformans* chs3Δ, as depletion of these cells fully rescues mice from death [13]. To delineate the specific effector mechanisms responsible for this immunopathology, we utilized a series of knockout mouse models to assess the roles of inflammatory cytokines, adaptive immune cells, and neutrophil-derived cytotoxic functions. We found that NADPH oxidase 2 (NOX2)-derived reactive oxygen species (ROS) were essential for the lethality observed following *chs3Δ* inoculation, whereas IL-6, myeloperoxidase (MPO), and adaptive immune cells were dispensable. Strikingly, NOX2-deficient mice exhibited preserved neutrophil recruitment to the lungs but displayed significantly blunted cytokine responses and complete survival without clinical signs of illness. In contrast, Rag1-and MPO-deficient mice succumbed with kinetics similar to wild-type controls. These findings demonstrate that neutrophils are not intrinsically pathogenic but become harmful through ROS-dependent amplification of inflammatory signaling. This work highlights ROS as a key effector of neutrophil-mediated immunopathology and suggests that targeting neutrophil effector functions—rather than neutrophils themselves—may offer a therapeutic strategy to mitigate hyperinflammatory responses, such as those observed in cryptococcal immune reconstitution inflammatory syndrome (IRIS)[14; 15; 16].

## MATERIAL AND METHODS

### Ethics Statement

All animal experiments were conducted in compliance with the Public Health Service Policy on Humane Care and Use of Laboratory Animals and adhered to the guidelines in the Guide for the Care and Use of Laboratory Animals. The study protocols were reviewed and approved by the Institutional Animal Care and Use Committee (IACUC) at the University of Tennessee Health Science Center (UTHSC) under protocol number 21-0268. All mice were housed in AALAC-accredited facilities located in the Regional Biocontainment Laboratory (RBL) at UTHSC under conditions of controlled temperature, humidity, and a 12-hour light/dark cycle. Mice were provided standard rodent chow and water ad libitum and were monitored daily for clinical signs of distress, including significant weight loss, labored breathing, or lethargy. Humane endpoints were applied as necessary to minimize suffering. The Laboratory Animal Care Unit (LACU) at UTHSC ensures compliance with all applicable provisions of the Animal Welfare Act, guidance from the Office of Laboratory Animal Welfare (OLAW), and the American Veterinary Medical Association Guidelines for the Euthanasia of Animals.

### Fungal strains and media

*C. neoformans* strain *chs3Δ* [13] was grown at 30°C, 300 rpm for 48 hours in 50 mL of YPD broth (1% yeast extract, 2% Bacto-peptone, and 2% dextrose). The cells were centrifuged, washed in endotoxin-free 1x PBS, and counted with a hemocytometer. For studies utilizing heat-killed organism, after diluting to the desired cell concentration in PBS, the inoculum was heated at 70°C for 15 minutes. Complete killing was assayed by plating for CFUs.

### Mice

C57BL/6 (000664), IL6^-/-^ (002650), Rag1^-/-^ (002216), MPO^-/-^ (004265), and gp91^phox–^ (002365) mice were obtained from Jackson Laboratory (Bar Harbor, ME). Mice were 6 to 8 weeks old at the time of inoculation, and all experiments were conducted with an equal mix of both male and female mice to avoid sex biases. All animal protocols were reviewed and approved by the Animal Studies Committee of the University of Tennessee Health Science Center and conducted according to National Institutes of Health guidelines for housing and care of laboratory animals.

### Pulmonary inoculations

Mice were anesthetized with isoflurane and inoculated by orotracheal aspiration with 1 × 10^7^ CFU of heat-killed organism in 50 μl of sterile PBS. The mice were fed *ad libitum* and monitored daily for symptoms. For survival studies, mice were sacrificed when their body weight fell below 80% of their weight at the time of inoculation. For cytokine analysis, flow cytometry studies, and histology, mice were euthanized at 4-, 6-, 8-, or 12-hours post-inoculation by CO_2_ inhalation, and the lungs were harvested.

### Histology

Mice were sacrificed according to approved protocols, perfused intracardially with sterile PBS, and the lungs inflated with 10% formalin. Lung tissue was then fixed for 48 hours in 10% formalin and submitted to HistoWiz Inc. (histowiz.com) for histology using a Standard Operating Procedure and fully automated workflow. Samples were processed, embedded in paraffin, sectioned at 4μm, and stained using hematoxylin-eosin (H&E). After staining, sections were dehydrated and film coverslipped using a TissueTek-Prisma and Coverslipper (SakuraUSA, Torrance, CA). Whole slide scanning (40x) was performed on an Aperio AT2 (Leica Biosystems, Wetzlar, Germany).

### Cytokine Analysis

Cytokine levels in lung tissues were analyzed using the Bio-Plex Protein Array System (Bio-Rad Laboratories, Hercules, CA). Briefly, lung tissue was excised and homogenized in 2 ml of ice-cold PBS containing 1X Pierce Protease Inhibitor cocktail (Thermo Scientific, Rockford, IL). After homogenization, Triton X-100 was added to a final concentration of 0.05%, and the samples were clarified by centrifugation. Supernatant fractions from the pulmonary homogenates were then assayed using the Bio-Plex Pro Mouse Cytokine 23-Plex (Bio-Rad Laboratories) for the presence of IL-1α, IL-1β, IL-2, IL-3, IL-4, IL-5, IL-6, IL-9, IL-10, IL-12 (p40), IL-12 (p70), IL-13, IL-17A, granulocyte colony stimulating factor (G-CSF), granulocyte monocyte colony stimulating factor (GM-CSF), interferon-γ (IFN-γ), CXCL1/keratinocyte-derived chemokine (KC), CCL2/monocyte chemotactic protein-1 (MCP-1), CCL3/macrophage inflammatory protein-1α (MIP-1α), CCL4/MIP-1β, CCL5/regulated upon activation, normal T cell expressed and secreted (RANTES) and tumor necrosis factor-α (TNF-α).

### Flow Cytometry

Cell populations in the lungs were identified by flow cytometry. Briefly, lungs from individual mice were enzymatically digested at 37°C for 30 min in digestion buffer (DMEM containing 0.05 mg/mL of Liberase TM and 0.02 mg/mL DNase I (Roche, Indianapolis, IN)). The digested tissues were then successively passed through sterile 70 and 40 µm pore nylon strainers (BD Biosciences, San Jose, CA). Erythrocytes in the strained suspension were lysed by incubation in NH_4_Cl buffer (0.859% NH_4_Cl, 0.1% KHCO_3_, 0.0372% Na_2_EDTA; pH 7.4; Sigma-Aldrich) for 3 min on ice, followed by the addition of a 2-fold excess of PBS. The leukocytes were then collected by centrifugation, resuspended in sterile PBS, and stained using eBioscience Fixable Viability Dye eFluor™ 506 (1:500; Invitrogen, Carlsbad, CA) for 30 min at 4° C in the dark. Following incubation, samples were washed and resuspended in FACS buffer (PBS, 0.1% BSA, 0.02% NaN_3_, 2 mM EDTA) and incubated with CD16/CD32 (Fc Block™; BD Biosciences, San Jose, CA) for 5 min. For flow cytometry, 1×10^6^ cells were incubated for 30 min at 4° C in the dark with optimal concentrations of fluorochrome-conjugated antibodies (Table S1 for antigen, clone, and source) diluted in Brilliant Stain Buffer (BD Biosciences). After three washes with FACS buffer, the cells were fixed in 2% ultrapure paraformaldehyde. For data acquisition, >200,000 events were collected on a NovoCyte 3000 flow cytometer (Agilent, Santa Clara, CA), and the data were analyzed with FlowJo V10 (TreeStar, Ashland, OR). The absolute number of cells in each leukocyte subset was determined by multiplying the absolute number of CD45^+^ cells by the percentage of cells stained by fluorochrome-labeled antibodies for each cell population analyzed.

### Bronchoalveolar lavage

BAL samples were obtained by performing one mL lavages with ice-cold PBS containing 2mM EDTA. The samples were then clarified by centrifugation, and the supernatants were treated with a protease inhibitor before being frozen at-80 °C. LTB4 was measured using an LTB4 ELISA kit (Cayman) according to the manufacturer’s protocol. Cytokines were analyzed using Mouse CXCL1/KC ELISA kit (R&D Systems), Mouse CXCL2/MIP-2α ELISA kit (R&D Systems), and Mouse C5a ELISA kit (R&D Systems).

### Statistics

Data were analyzed using GraphPad Prism, version 10.0 (GraphPad Software, Inc., La Jolla, CA). The one-way analysis of variance (ANOVA) with the Tukey’s multiple-correction test was used to compare more than two groups. Kaplan-Meier survival curves were compared using the Mantel-Cox log rank test. p values <0.05 were considered significant.

## RESULTS

### IL-6 and adaptive immunity contribute minimally to *chs3Δ*-induced mortality

We have previously shown that inoculation with a heat-killed chitosan-deficient strain of *Cryptococcus neoformans* (*chs3Δ*) results in rapid and neutrophil-dependent mortality. To determine whether specific cytokines contribute to this lethal inflammatory response, we focused on interleukin-6 (IL-6), which plays a critical role in neutrophil recruitment and was the most abundantly expressed cytokine in the lungs of the c*hs3Δ-*inoculated mice. Wild-type (WT) and IL-6 knockout (IL-6^-/-^) mice were inoculated with 10^7^ heat-killed (HK) *chs3Δ* yeast and monitored for survival. We observed that there was a slight increase in the survival of the IL-6^-/-^ mice compared to WT, with a median survival of 2 days compared to 1 day for the WT mice (Figure 1A). Although this difference was statistically significant, it was modest, suggesting that the high levels of IL-6 observed may only play a minor role in the mortality of *chs3Δ-*inoculated mice. These data indicate that IL-6 is not the primary driver of immunopathology or neutrophil recruitment in this model.

**Figure 1.**
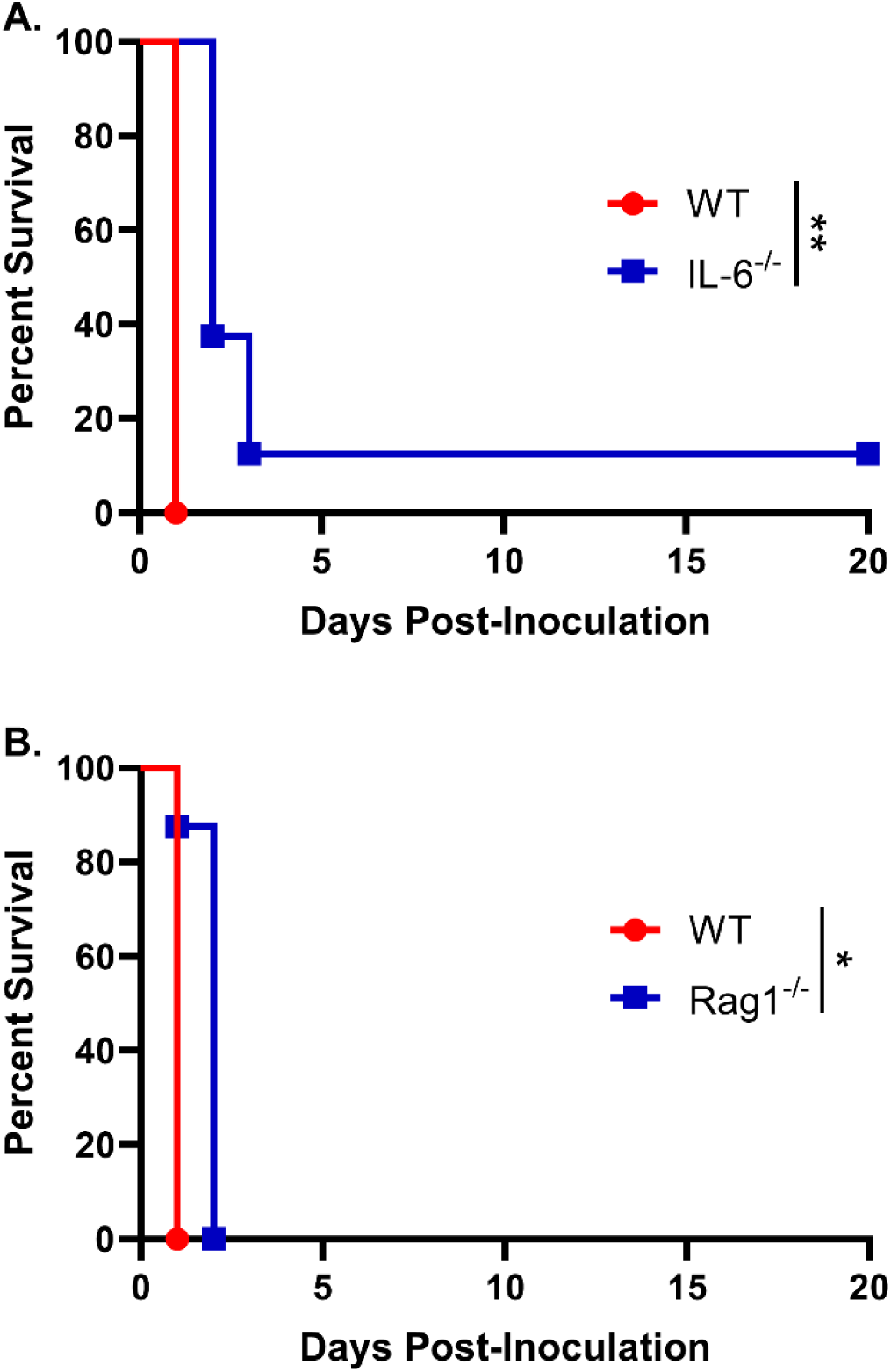
IL-6 and adaptive immunity contribute minimally to *chs3Δ*-induced mortality: Male and female A) C57BL/6 and IL-6^-/-^ mice or B) C57BL/6 and Rag1^-/-^ mice were inoculated with 10^7^ CFUs of HK *chs3Δ* and monitored for survival. Survival of the animals was recorded as mortality of mice for 20 days post inoculation. Data are representative of one experiment with 8 mice per group. Virulence was determined using Mantel-Cox curve comparison, with statistical significance determined by the log rank test (*, *P* <0.05, **, *P* <0.005).

As the patient population susceptible to cryptococcosis often includes individuals with compromised adaptive immunity, particularly CD4⁺ T cell deficiencies, we next assessed whether T and B cells contribute to the host response to *chs3Δ*. To this end, WT and Rag1^-/-^ mice, which lack mature T and B lymphocytes [17], were inoculated with 10^7^ HK *chs3Δ* yeast and monitored for survival. We observed that Rag1^-/-^ mice exhibited a modest increase in survival compared to WT controls, with Rag1^-/-^ mice succumbing on day 2 post-inoculation, whereas WT mice succumbed by day 1 (Figure 1B). Similar to IL-6^-/-^ mice, this modest delay in mortality suggests that the adaptive immune response plays only a minor role in the *chs3Δ*-induced lethality. These results support the conclusion that the rapid mortality observed in this model is primarily driven by innate immune mechanisms, particularly those involving neutrophils.

### NOX2-dependent ROS, but not MPO, mediate neutrophil-dependent lethality

Inoculation with HK *chs3Δ* induces a rapid and robust neutrophilic response, and previous studies have demonstrated that neutrophil depletion rescues mice from the associated mortality [13], implicating neutrophils as central mediators of the observed immunopathology. However, the precise neutrophil effector mechanisms responsible for this lethality remain unclear. Neutrophils deploy several antimicrobial effectors, including neutrophil elastase, myeloperoxidase (MPO), and reactive oxygen species (ROS) generated by NADPH oxidase 2 (NOX2), which can also damage host tissues.

To define the contributions of these effectors, we evaluated the responses of MPO knockout (MPO^-/-^) and NOX2-knockout (gp91^phox–^) mice. Male and female MPO^-/-^, gp91^phox–^, and WT mice were inoculated with 10^7^ HK *chs3Δ* yeast and monitored for survival. All MPO^-/-^ mice succumbed with the same rapid kinetics as WT mice (Figure 2A), indicating that MPO-derived hypochlorous acid is not responsible for the lethal phenotype.

**Figure 2.**
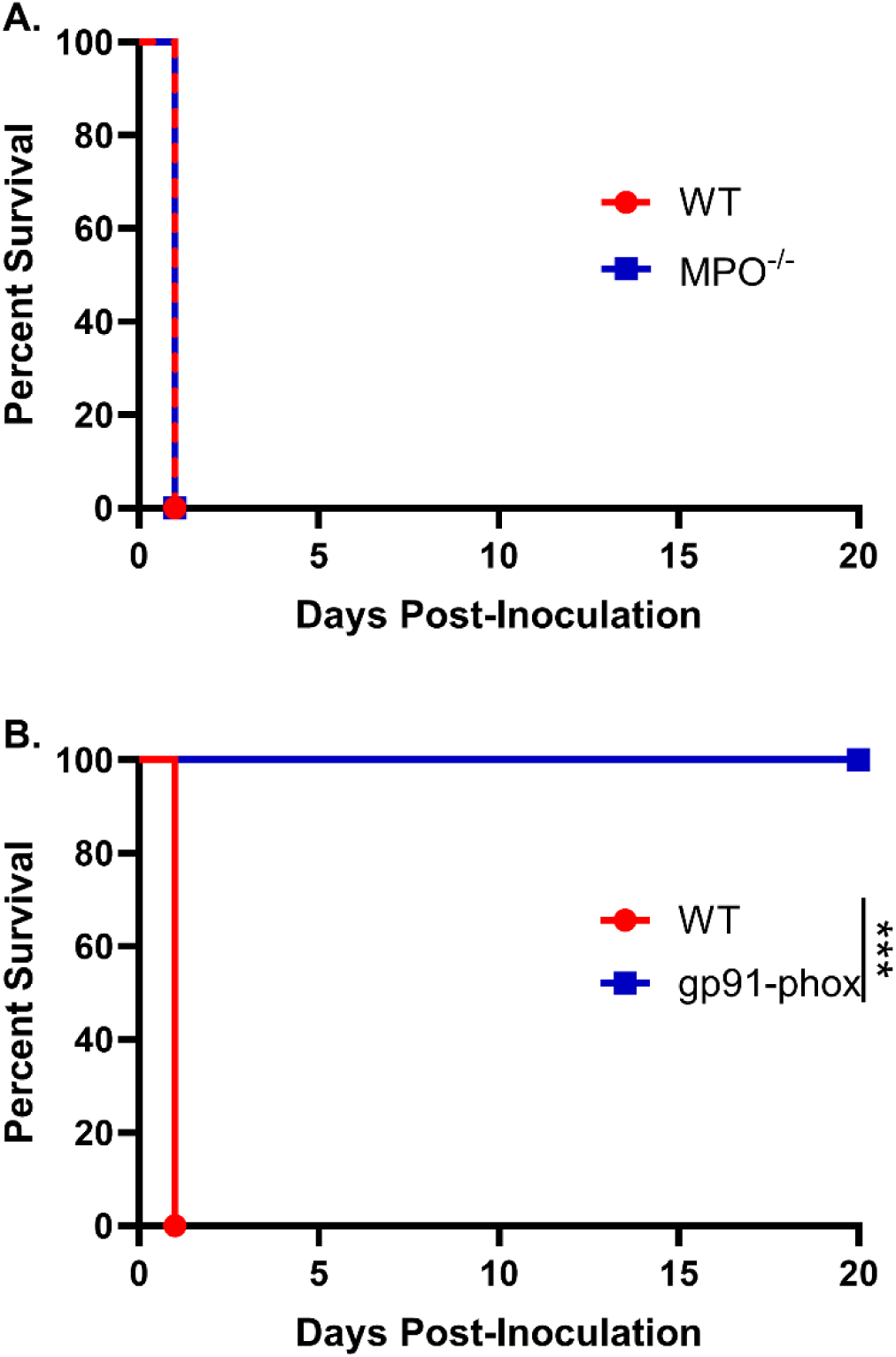
NOX2-dependent ROS, but not MPO, mediate neutrophil-dependent lethality: Male and female A) C57BL/6 and MPO^-/-^ mice or B) C57BL/6 and gp91^phox-^ mice were inoculated with 10^7^ CFUs of HK *chs3Δ* and monitored for survival. Survival of the animals was recorded as mortality of mice for 20 days post inoculation. Data are representative of one experiment with A) 8 mice per group or B) two experiments with a total of 12 mice per group. Virulence was determined using Mantel-Cox curve comparison, with statistical significance determined by the log rank test (***, *P* <0.001).

In striking contrast, gp91^phox–^ mice were completely protected from *chs3Δ*-induced mortality. All gp91^phox–^ mice survived the inoculation and exhibited no signs of morbidity, maintained normal grooming behavior throughout the observation period (Figure 2B). This is in stark contrast to the neutrophil-depleted mice, which while also surviving exhibited transient illness marked by ruffled fur and weight loss before eventually recovering [13]. These results demonstrate that NOX2-derived ROS, and not neutrophils per se, are the critical drivers of the immunopathology associated with *chs3Δ* inoculation.

Moreover, the absence of morbidity in gp91^phox–^ mice suggests that neutrophils lacking ROS production may play a non-pathogenic or potentially regulatory role during infection.

### NOX2-deficient mice exhibit attenuated cytokine responses despite robust neutrophil infiltration

Given that NOX2-deficient mice were completely protected from *chs3Δ*-induced lethality, we next sought to characterize the pulmonary immune response in these animals. To assess local cytokine production, WT and gp91^phox–^ mice were inoculated with 10^7^ HK *chs3Δ* yeast. At 4-, 8-, and 12-hours post-inoculation, lungs were harvested, homogenized, and cytokine/chemokine responses were determined from the lung homogenates using the Bio-Plex protein array system. We observed an increase in multiple cytokines (Figure. S1). As expected, WT mice displayed significantly elevated levels of IL-6, G-CSF, and CXCL1/KC, with cytokine levels peaking at 8 hours and remaining elevated at 12 hours post-inoculation (Figure 3). In contrast, gp91^phox–^ mice exhibited markedly reduced levels of all three cytokines, with concentrations approaching baseline by 12 hours (Figure 3). These findings suggest that NOX2-dependent ROS are required for the amplification or maintenance of the proinflammatory cytokine milieu in the lung following *chs3Δ* challenge.

**Figure 3.**
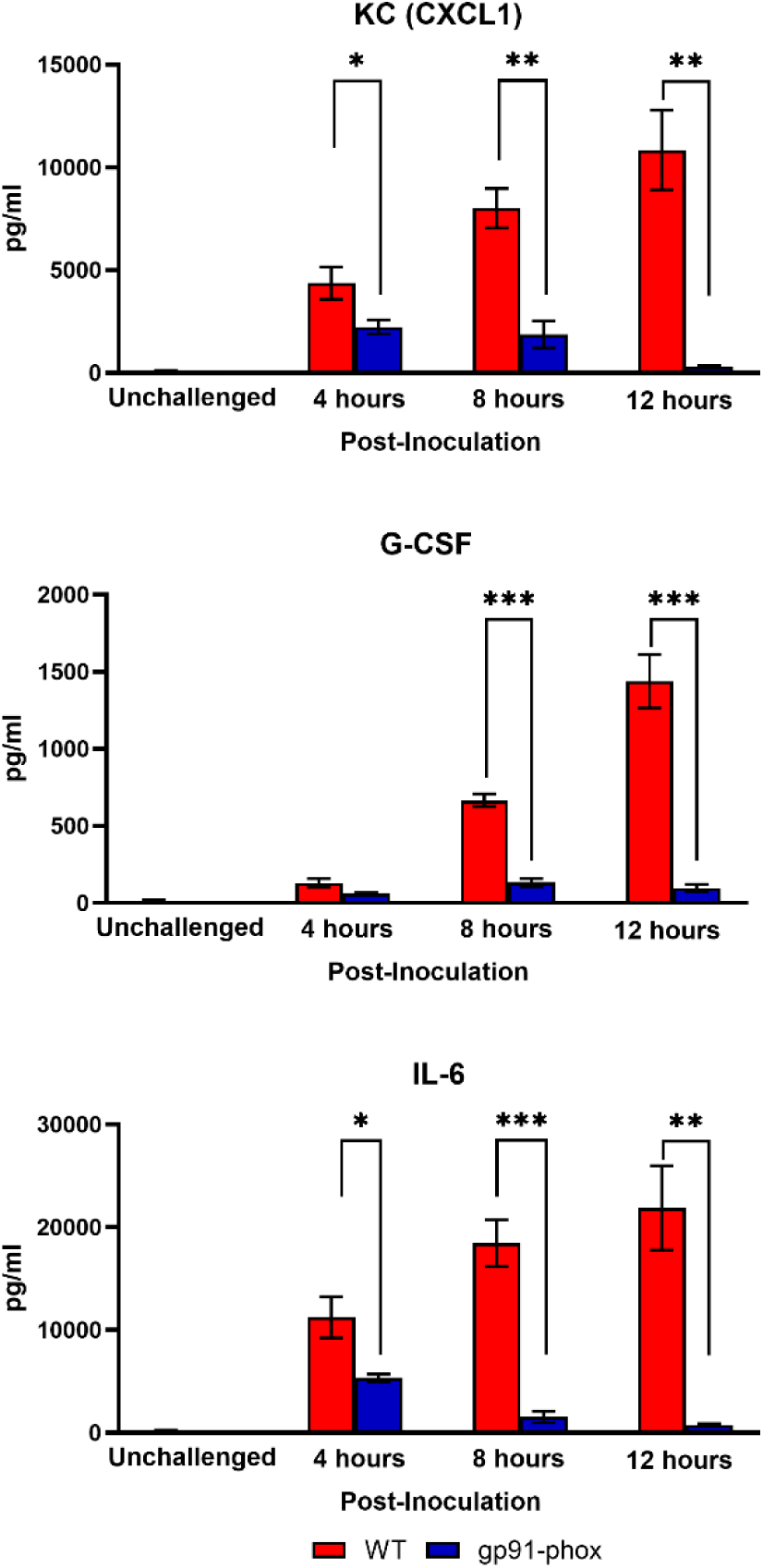
**NOX2 KO mice have reduced cytokine and chemokine levels**. Male and female C57BL/6 or gp91^phox-^ mice were inoculated with 10^7^ CFUs of HK *chs3Δ*. Homogenates were prepared from the lungs of each group at the indicated time point as well as a phosphate-buffered saline (PBS) control for each group. Cytokine/chemokine responses were determined from the lung homogenates. Data are cumulative of two experiments for a total of 8 mice per group per timepoint. Values are means ± standard errors of the means (SEM). (*, *P* <0.05, **, *P* <0.005, ***, *P* <0.001).

Because NOX2-deficient mice survived *chs3Δ* inoculation and had significantly reduced chemokines, we hypothesized that the mice lived because there was no or reduced neutrophil recruitment. Despite the dampened cytokine response, histological examination of lung tissues revealed no overt difference in cellular infiltration between WT and gp91^phox–^ mice. At 12 hours post-inoculation, both groups displayed numerous inflammatory foci diffusely distributed throughout the lung parenchyma, composed predominantly of granulocytic cells (Figure 4A). This result was unexpected given the suppression of chemokine signals in gp91^phox–^ mice.

**Figure 4.**
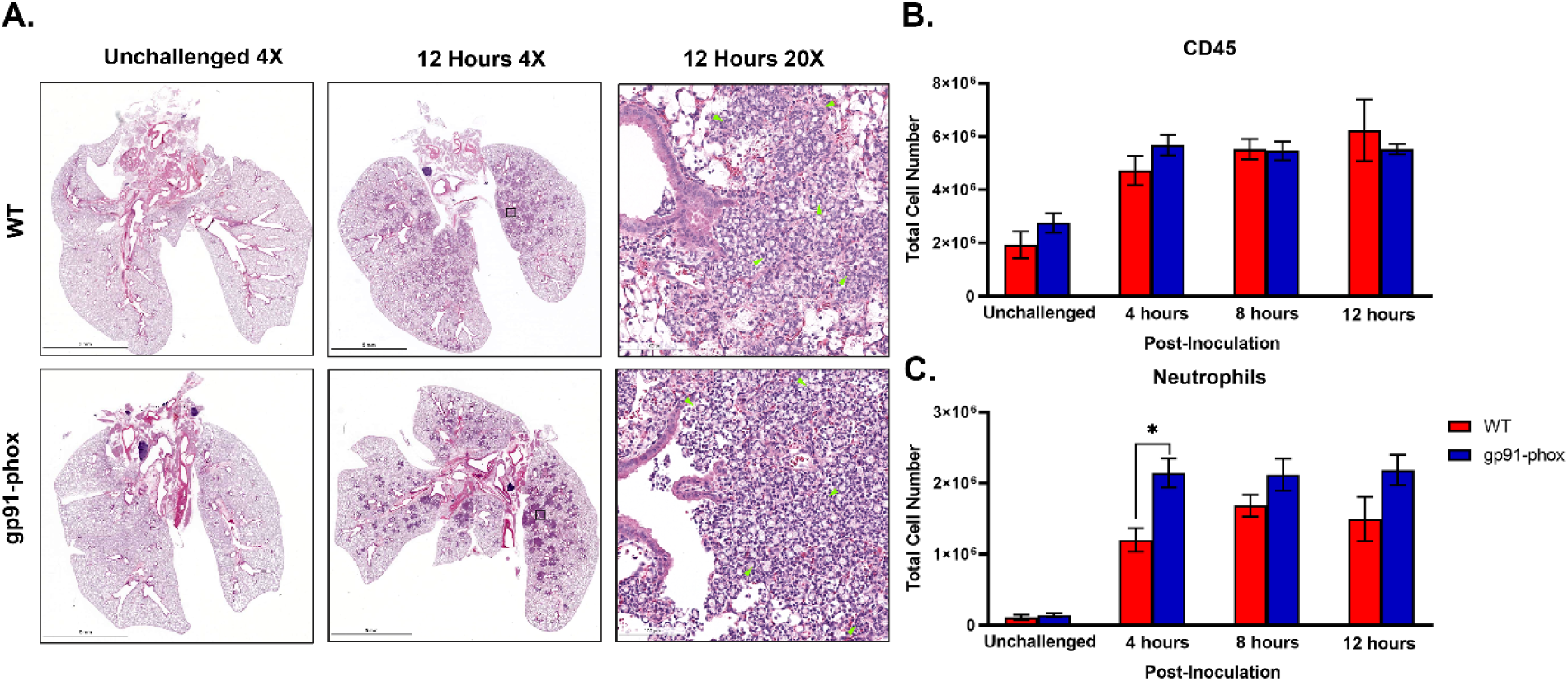
**NOX2 KO mice have similar amounts of neutrophils:**. A) Male and female C57BL/6 or gp91^phox-^ mice were inoculated with 10^7^ CFUs of HK *chs3Δ*. At 12 h post-inoculation, the lungs were harvested, embedded, sectioned, and processed for hematoxylin and eosin staining. Green arrowheads demarcate neutrophils. Images are representative of two independent experiments using three mice per group. B,C) At the indicated time point, pulmonary leukocytes were isolated from the lungs of mice of each group and subjected to flow cytometry analysis. Data are cumulative of two experiments for a total of 8 mice per group per timepoint. Values are means ± standard errors of the means (SEM). (*, *P* <0.05). Neutrophils are defined as CD11b^+^/CD24^+^/Ly6G^+^/CD45^+^.

To quantitatively assess leukocyte recruitment, we performed flow cytometry on lung leukocytes isolated from WT and gp91^phox–^ mice inoculated with 10^7^ HK *chs3Δ* yeast at 4-, 8-, and 12-hours post-inoculation. Consistent with the above histology data, total CD45⁺ leukocyte counts were comparable between WT and gp91^phox-^ groups at all time points (Figure 4B). Notably, gp91^phox-^ mice exhibited slightly increased numbers of neutrophils at 4 hours post-inoculation, which remained stable through 12 hours, whereas WT mice showed a gradual increase over time (Figure 4C). We also observe a significant decrease in Ly6C^hi^ monocytes and interstitial macrophages in the gp91^phox-^ mice 12-hours post-inoculation, likely due to the reduction in inflammatory cytokines (Figure. S2). These results indicate that neutrophil recruitment occurs independently of NOX2 and is not diminished in the absence of ROS.

Taken together, these data reveal that while NOX2-deficient mice mount a robust neutrophilic response to *chs3Δ*, the associated cytokine storm is abrogated, and the animals are protected from lethal immunopathology. These findings suggest that neutrophil recruitment alone is insufficient to cause host damage in this model and that NOX2-derived ROS are essential for driving both the proinflammatory cytokine response and the resulting pathology

### Differential regulation of CXCR2 ligands may drive neutrophil recruitment in the absence of ROS

Despite the significantly reduced levels of CXCL1/KC and G-CSF in NOX2-deficient mice, we observed robust neutrophil recruitment to the lungs following *chs3Δ* inoculation (Figure 4). Multiple mediators can induce neutrophil recruitment to the site of infection. To determine which mediators might support this ROS-independent neutrophil influx, we analyzed bronchoalveolar lavage fluid (BALF) from WT and gp91^phox-^ mice at 12 hours post-inoculation for common neutrophil chemoattractants.

In WT mice, high concentrations of leukotriene B4 (LTB4) and complement component C5a, two well-established neutrophil chemoattractants, were detected. In contrast, both LTB4 and C5a levels were significantly diminished in gp91^phox-^ mice (Figure 5A & 5B), ruling out their contribution to the preserved neutrophil infiltration in the absence of ROS. We next assessed expression of the CXCR2 ligands CXCL1/KC and CXCL2/MIP-2α. As previously reported, CXCL1/KC was highly expressed in WT mice but was dramatically reduced in gp91^phox-^ animals (Figure 5C). Conversely, CXCL2/MIP-2α levels were significantly elevated in NOX2-deficient mice compared to WT controls (Figure 5D). This inverse regulation of CXCL1/KC and CXCL2/MIP-2α suggests that, in the absence of ROS, neutrophil recruitment may be maintained through an alternative CXCL2/MIP-2α-driven pathway. Given that both chemokines signal through CXCR2 [18], these findings imply that neutrophil migration in gp91^phox-^mice is not impaired but rather redirected via differential chemokine signaling. Together, these data point to a ROS-independent but CXCR2-dependent mechanism of neutrophil recruitment in response to *chs3Δ* challenge.

**Figure 5.**
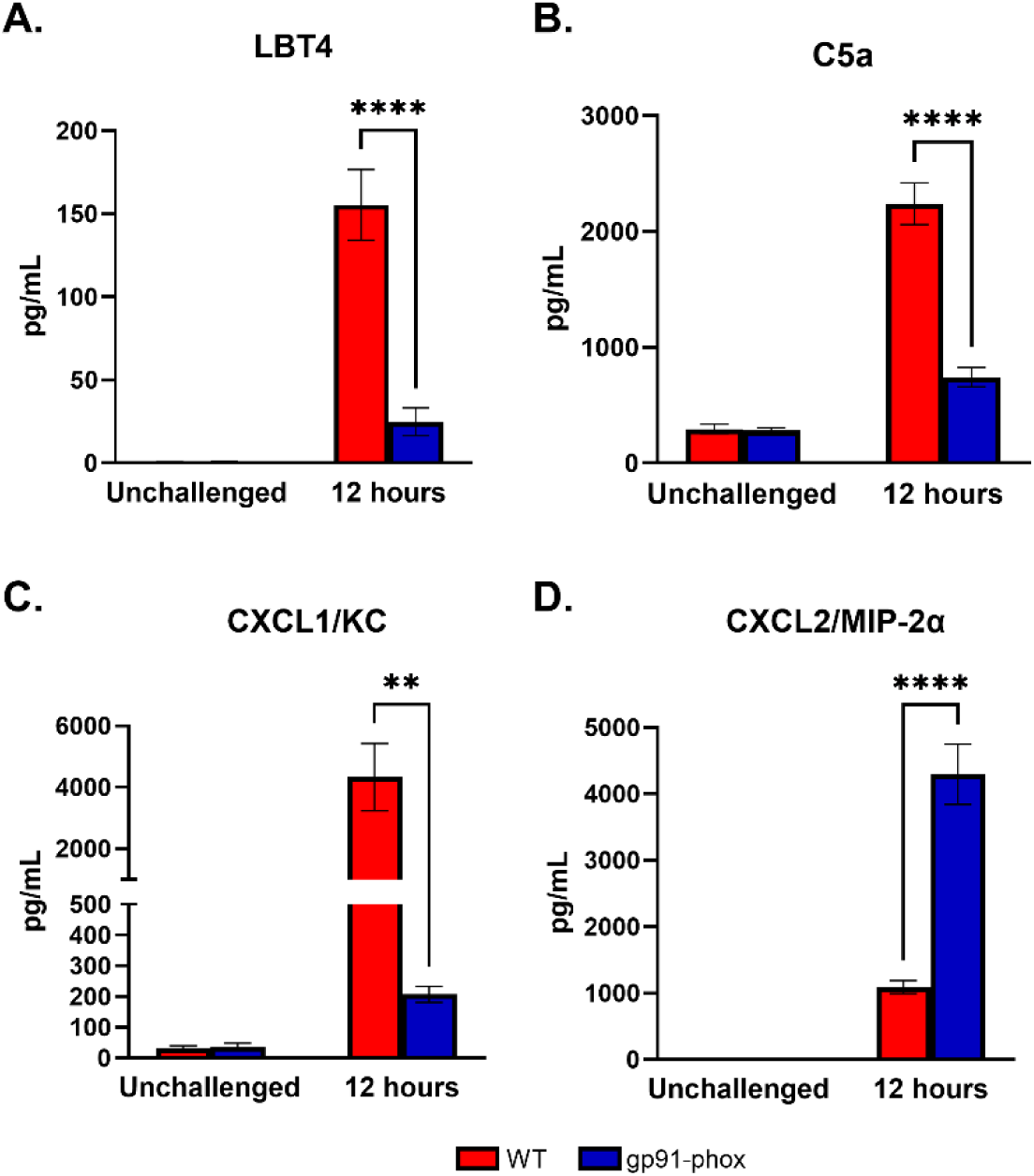
Differential regulation of CXCR2 ligands may drive neutrophil recruitment in the absence of ROS: Male and female C57BL/6 or gp91^phox-^ mice were inoculated with 10^7^ CFUs of HK *chs3Δ*. BAL was collected from each group at 12 h post-inoculation, as well as a phosphate-buffered saline (PBS) control for each group. Data are cumulative of two experiments for a total of 8 mice per group per timepoint. Values are means ± standard errors of the means (SEM). (**, *P* <0.005, ***, *P* <0.001).

## DISCUSSION

Our findings reveal a critical and previously unrecognized role for neutrophil-derived reactive oxygen species (ROS) in driving the aberrant immune response elicited by a chitosan-deficient *C. neoformans chs3Δ* mutant. In contrast to the expected protective function of ROS in fungal infections, we demonstrate that NADPH oxidase 2 (NOX2)-dependent ROS production is essential for the mortality observed in mice inoculated with heat-killed *chs3Δ* cells. Notably, NOX2-deficient mice were completely protected from death, despite maintaining robust pulmonary neutrophil recruitment. These data provide compelling evidence that ROS are the primary mediators of immunopathology in this model, and that neutrophils, in the absence of ROS, do not contribute to the damaging inflammatory response.

This represents a significant divergence from canonical models in which ROS are required for fungal clearance and host protection. In *Aspergillus fumigatus* infection, for example, NOX2 deficiency increases fungal susceptibility and exacerbates invasive disease [19; 20; 21]. Similarly, patients with chronic granulomatous disease (CGD), who lack functional NADPH oxidase, are highly susceptible to invasive fungal infections [22; 23]. However, these protective roles of ROS may not extend to all fungal pathogens or all infection contexts. For instance, in *C. deneoformans* infection, NOX2-deficient mice had lower fungal burdens and an enhanced Th1 response, although early inflammatory kinetics and neutrophil involvement were not assessed [24]. Our data now expands these findings by highlighting a specific context, chitosan deficiency, where ROS play a pathological rather than protective role.

The *chs3Δ* strain triggers a highly neutrophilic inflammatory response characterized by elevated lung IL-6, G-CSF, and CXCL1/KC [13]. Our current data demonstrate that these cytokines are significantly reduced in NOX2-deficient mice, suggesting that ROS plays a role in amplifying cytokine production. This finding is consistent with previous studies that have shown ROS can act as signaling molecules to promote the expression of proinflammatory cytokines through the NF-κB and MAPK pathways [25]. However, the neutrophil response in NOX2-deficient mice was not attenuated. In fact, early recruitment was even more pronounced, suggesting that neutrophil migration into the lung can be uncoupled from ROS-mediated cytokine signaling.

This led us to investigate alternative neutrophil chemoattractants. Although CXCL1/KC and G-CSF were significantly diminished in NOX2-deficient mice, CXCL2/MIP-2α was paradoxically elevated. Both CXCL1/KC and CXCL2/MIP-2α bind CXCR2, but they are produced by different cell types and are differentially regulated [18]. CXCL1/KC is mainly derived from epithelial and endothelial cells, while neutrophils themselves often produce CXCL2/MIP-2α and can be part of an autocrine amplification loop [18; 26]. Both CXCL1/KC and CXCL2/MIP-2α have been observed in cryptococcal infections [27; 28; 29]. Our data suggest a shift from CXCL1/KC-to CXCL2/MIP-2α-dominated chemotaxis in NOX2-deficient mice, with potential implications for the nature of the recruited neutrophil subsets.

Neutrophil heterogeneity has been increasingly recognized as a key factor in shaping immune responses during infection, cancer, and sterile inflammation [30; 31; 32]. One marker that distinguishes neutrophil subsets is CD101, a surface glycoprotein associated with mature neutrophils. Immature CD101⁻ neutrophils exhibit enhanced pro-inflammatory transcriptional signatures and have been implicated in tissue-damaging responses [33; 34]. In zymosan-induced inflammation, NOX2-deficient mice mobilize an increased proportion of CD101⁻ neutrophils to the lung, and these cells display a distinct and more inflammatory gene expression profile compared to their WT counterparts [35].

Additionally, during cryptococcal infection, distinct neutrophil subsets with either oxidative stress-associated or cytokine-producing transcriptional signatures have been described [36]. Our flow cytometry data confirm high neutrophil numbers in the lungs of *chs3Δ*-inoculated gp91^phox-^ mice; however, these animals remain asymptomatic. Whether this discrepancy is due to reduced effector potential or a more tolerogenic neutrophil phenotype is under investigation. It is possible that NOX2 deficiency skews neutrophil differentiation toward a less inflammatory phenotype or alters their activation state in a way that preserves tissue integrity. This may also explain why NOX2-deficient neutrophils, though abundant, do not drive pathology. Given the accumulating evidence that phenotypic heterogeneity among neutrophils can shape the immune outcome, further profiling of neutrophil subsets in this model is warranted.

The finding that neutrophils are not inherently pathogenic in the absence of ROS is particularly relevant for conditions such as cryptococcal immune reconstitution inflammatory syndrome (IRIS), which is marked by exaggerated immune responses following antiretroviral therapy (ART) in HIV-positive individuals. IRIS is associated with increased neutrophilic inflammation and high levels of IL-6 and G-CSF [14; 15]. Our model recapitulates many features of IRIS, including neutrophil-dependent mortality in the absence of viable fungi, and identifies ROS as a potential therapeutic target. Notably, unlike neutrophil depletion, which prevents death but does not fully eliminate inflammation or weight loss [13], NOX2 deficiency protects without apparent morbidity, suggesting that selective targeting of neutrophil effector functions may offer clinical benefit without impairing host defense.

We also investigated other neutrophil-derived effectors, including myeloperoxidase (MPO), but found that MPO-deficient mice were not protected from *chs3Δ*-induced mortality. This indicates that ROS, and not MPO-generated hypochlorous acid, are the dominant cytotoxic species in this model. This finding is consistent with literature showing that while MPO contributes to pathogen killing, its role in inflammation-driven tissue damage is often redundant with ROS [22].

Our data also raise questions about upstream regulators of ROS-dependent pathology. Previous studies have identified leukotriene B4 (LTB4) and complement component C5a as potent neutrophil chemoattractants and activators in fungal infections [37; 38]. While both were elevated in WT mice, they were absent in NOX2-deficient mice, suggesting a ROS-dependent positive feedback loop. However, their absence did not prevent neutrophil accumulation, implying that they are not essential for neutrophil recruitment in this context, but may modulate activation state or effector functions.

Finally, this study underscores the complex interplay between fungal cell wall composition and host immunity. The *chs3Δ* strain lacks chitosan, exposing immunogenic cell wall components and triggering a profound immune response. The absence of ROS dampens this response at the cytokine level but not at the level of cell recruitment. Future work will aim to identify which fungal or host molecules act as ROS-dependent immune amplifiers, and how neutrophil subsets are shaped by ROS signaling.

## ACKNOWLEDGMENTS

This work was supported by a National Institutes of Health grant K22 AI148724, the Center of Pediatric Experimental Therapeutics, and start-up funds from the Department of Clinical Pharmacy and Translational Science. The funders had no role in study design, data collection, interpretation, or the decision to submit the work for publication.

---

**Supplementary Figure 1.**
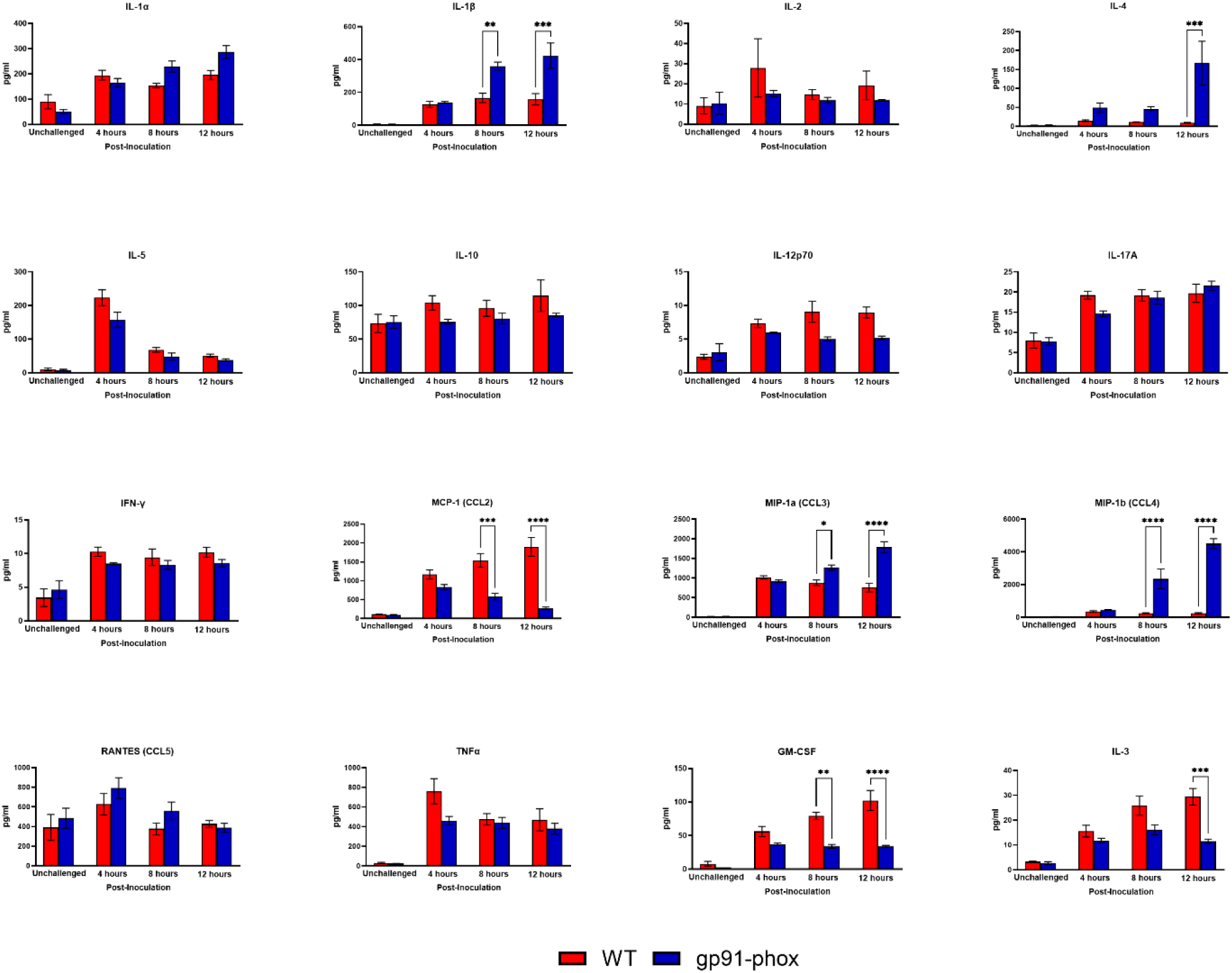
Cytokine/chemokine analysis: Male and female C57BL/6 or gp91^phox-^ mice were inoculated with 10^7^ CFUs of HK *chs3Δ*. Homogenates were prepared from the lungs of each group at the indicated time point as well as a phosphate-buffered saline (PBS) control for each group. Cytokine/chemokine responses were determined from the lung homogenates. Data are cumulative of two experiments for a total of 8 mice per group per timepoint. Values are means ± standard errors of the means (SEM). (*, *P* <0.05, **, *P* <0.005, ***, *P* <0.001).

**Supplementary Figure 2.**
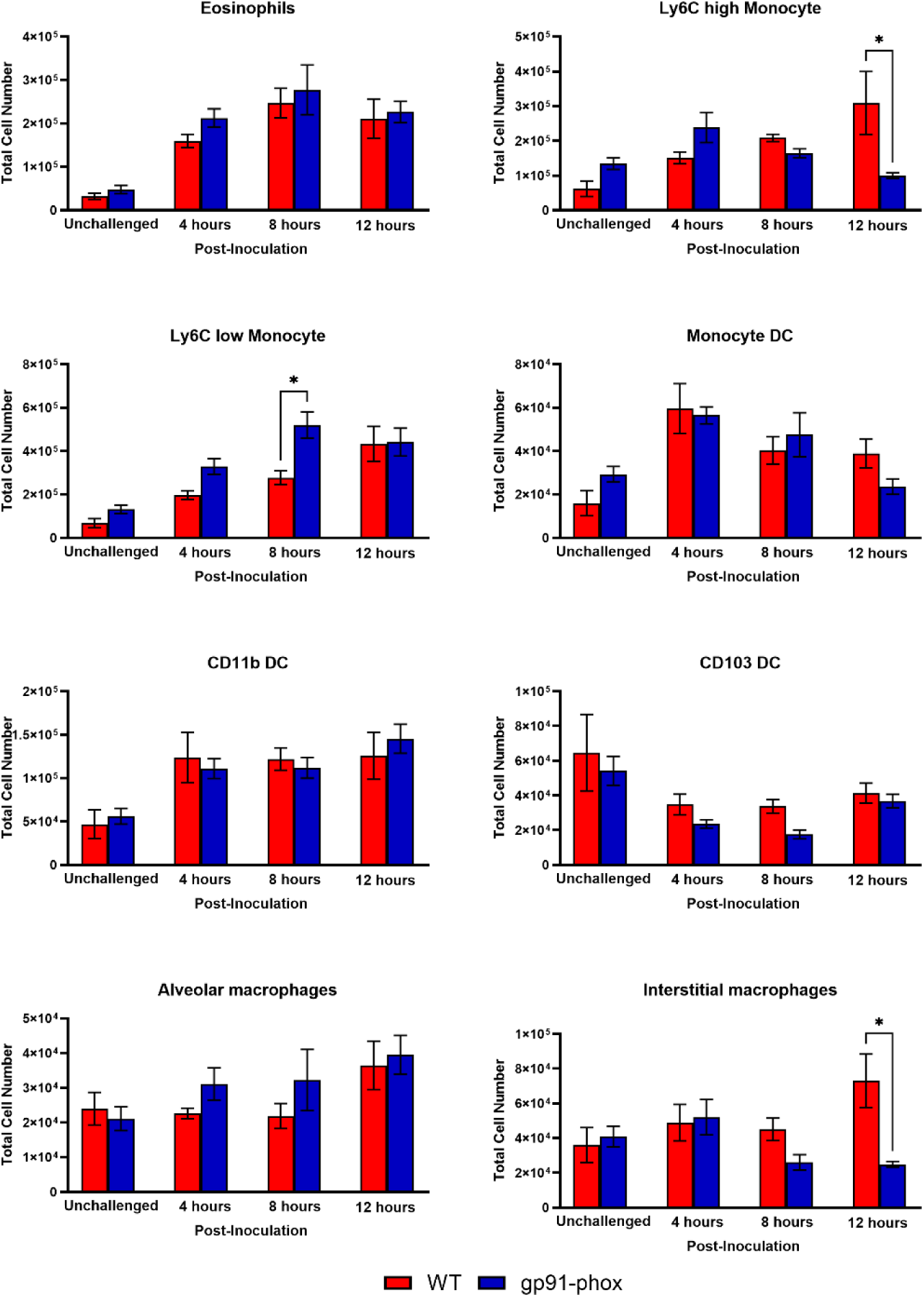
Flow cytometry analysis: Male and female C57BL/6 or gp91^phox-^ mice were inoculated with 10^7^ CFUs of HK *chs3Δ*. At the indicated time point, pulmonary leukocytes were isolated from the lungs of mice of each group and subjected to flow cytometry analysis. Data are cumulative of two experiments for a total of 8 mice per group per timepoint. Values are means ± standard errors of the means (SEM). (*, *P* <0.05).

**Table S1.**
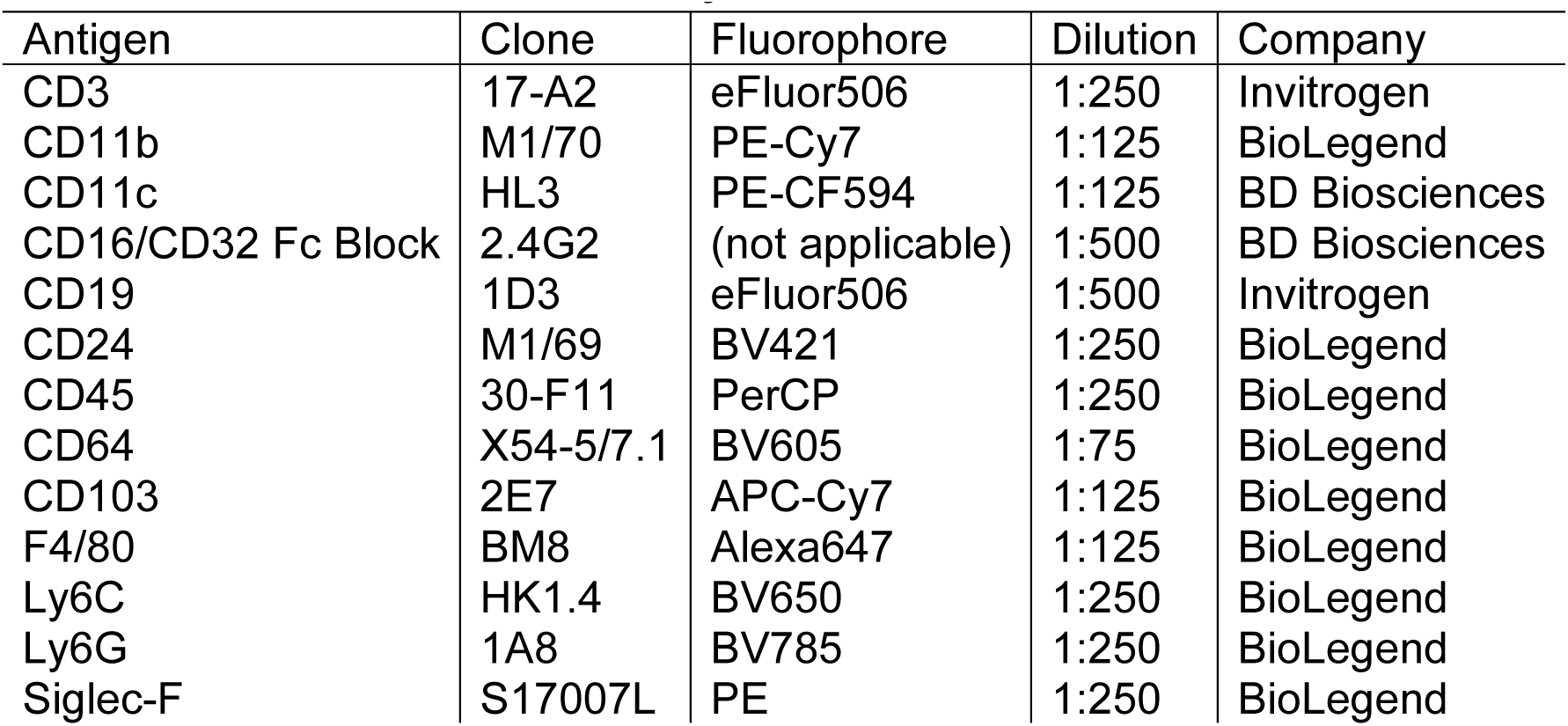
Antibodies for flow analysis.

